# Accelerating Minimap2 for accurate long read alignment on GPUs

**DOI:** 10.1101/2022.03.09.483575

**Authors:** Harisankar Sadasivan, Milos Maric, Eric Dawson, Vishanth Iyer, Johnny Israeli, Satish Narayanasamy

## Abstract

Long read sequencing technology is becoming increasingly popular for Precision Medicine applications like Whole Genome Sequencing (WGS) and microbial abundance estimation. Minimap2 is the state-of-the-art aligner and mapper used by the leading long read sequencing technologies, today. However, Minimap2 on CPUs is very slow for long noisy reads. ∼60-70% of the run-time on a CPU comes from the highly sequential chaining step in Minimap2. On the other hand, most Point-of-Care computational workflows in long read sequencing use Graphics Processing Units (GPUs). We present minimap2-accelerated (`mm2-ax`), a heterogeneous design for sequence mapping and alignment where minimap2’s compute intensive chaining step is sped up on the GPU and demonstrate its time and cost benefits.

We extract better intra-read parallelism from chaining without loosing mapping accuracy by forward transforming Minimap2’s chaining algorithm. Moreover, we better utilize the high memory available on modern cloud instances apart from better workload balancing, data locality and minimal branch divergence on the GPU. We show `mm2-ax` on an NVIDIA A100 GPU improves the chaining step with 5.41 - 2.57X speedup and 4.07 - 1.93X speedup : costup over the fastest version of Minimap2, `mm2-fast`, benchmarked on a Google Cloud Platform instance of 30 SIMD cores.

## Introduction

Long read sequencing is gaining more popularity with improved raw read accuracy, reduced end-to-end sequencing times, lower costs of adoption and ease of portability^1, 2^. Longer reads help span highly repetitive regions in the genome which short reads cannot. This helps applications like denovo assembly and structural variant calling^1, 3^. A recent study used long read sequencing to showcase the world’s fastest blood to variants workflow for genetic diagnosis at the point-of-care^2^. This further underlines the emerging significance of long read sequencing.

Amongst the many post-sequencing steps in long read processing workflows, sequence mapping and alignment is one of the first and amongst the most time and cost consuming steps. Sequence alignment^4^ in bioinformatics is a way of arranging the primary sequences of DNA, RNA or protein to identify regions of similarity while sequence mapping is a subset of alignment and only finds the approximate origin of query sequence in the target. We observe that sequence mapping and alignment is slow and users often spend costly cloud instance hours to keep up with high throughput sequencers^2^. This problem can worsen as the focus shifts to longer reads.

Additionally, we find that General Purpose Graphics Processing Units (GPGPUs or simply GPUs) are becoming increasingly popular for genomics processing. Several high throughput sequencers from Oxford Nanopore^5^ (GridION and PromethION series), Thermofisher’s Ion Proton 48^6^ and MGI’s DNBSEQ-T7^7^ have in-built GPUs. Many popular genome sequencing workflows also utilize GPUs for computation^8–10^.

In this work, we present minimap2-accelerated (`mm2-a`*x*) which speeds up minimap2 (`mm2`) on the GPU without loosing mapping accuracy and demonstrate its time and cost benefits.

## Background

### Minimap2: A brief overview

Minimap2 (`mm2`)^11^ is the state-of-the-art DNA/mRNA sequence mapper and aligner for the most popular long read sequencing platforms like Oxford Nanopore Technologies (ONT) and Pacific Biosciences (PacBio)^12^. While BLAST^4^ (using seed-extend paradigm) remains a powerful tool for full genome alignment, it is very slow especially on very long reads. For faster alignments, more recent aligners^13–17^ including `mm2` filter seeds prior to the final step of base-level alignment. `mm2`’s algorithm is based on the seed-chain-align paradigm (detailed in Fig. 1) and has an offline pre-processing step to build index from target reference. In the offline pre-processing step, the reference genome is indexed to a multimap using a hash table with the popular time and space-saving k-mer samples called minimizers^18^ as the key and minimizer locations on the reference as the values.

**Figure 1.**
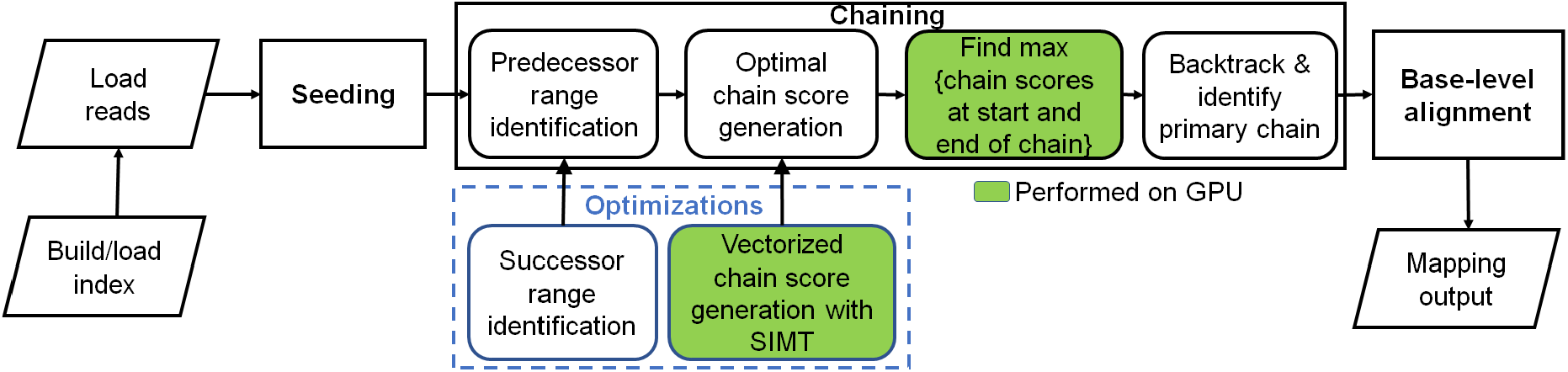
Minimap2 operates in 3 main steps: seeding, chaining and base-level alignment. Our optimizations to chaining are shown in blue box. Boxes with green fill show chaining sub-tasks which we perform on the GPU instead of CPU.

Seeding is fast and identifies short fixed-length exact matches (minimizer seeds) between a read and a reference sequence. When `mm2` processes a sequenced read, minimizers from the read are used to query the reference index for exact matches (anchors). These anchors are then sorted based on position in the reference and then passed onto the next step, chaining.

Chaining takes anchors sorted based on position in the reference as the input and identifies collinear ordered sub-sets of anchors called chains such that no anchor is used in more than one chain. `mm2` implements chaining via 1-dimensional dynamic programming^19^ where a complex problem is recursively broken down into simpler sub-problems. In summary, chaining sub-selects a few regions (chains) on the target reference and reduces the work for the next step of base-level alignment.

Further, if base-level alignment is requested, a 2-dimensional dynamic programming (Needleman-Wunsch^20^ with Suzuki-Kazahara formulation^21^) is applied to extend from the ends of chains in order to close the gaps between adjacent anchors in the chains.

`mm2` is considered accurate and has multiple use cases^11^. It may be used to map long noisy DNA/cDNA/mRNA reads, short accurate genomics reads, to find overlaps between long reads and for aligning with respect to a full reference genome or genome assembly. It is only for full genome or assembly alignment that `mm2` proceeds from chaining to the last step of base level alignment.

For a more detailed understanding of how seeding and base-level alignment operates, one may refer to prior literature^11, 22^. In the context of this work, we discuss chaining in-depth as it is the bottleneck stage in `mm2` we optimize.

### Minimap2: Sequential chaining

Chaining is the second step in `mm2` and sub-selects regions on the target reference where the last step of base-level alignment may be performed. An anchor is a short exact-match on the reference and is a 3-tuple (coordinate on the reference sequence, coordinate on the query sequence, length of the anchor). Chaining performs 1-dimensional dynamic programming on the input sorted anchors (from the seeding step) to identify collinear ordered sub-sets of anchors called chains such that no anchor is used in more than one chain. The chaining task can further be sub-divided into 4 sub-tasks: predecessor range selection, optimal chain score generation, finding maximum score from start and end of chains, and backtracking and primary chain identification.

Predecessor range selection is performed for every anchor in the output sorted list of anchors from the seeding step in order to dynamically calculate the number of preceding anchors (0-5000) to which chaining is attempted. While Guo et al.^23^ chose a static predecessor range of 64 for every anchor, `mm2` does a dynamic calculation of the predecessor range by finding all predecessors within a distance threshold.

Optimal chain score generation finds the preceding anchor within the predecessor range which yields the maximum chain score, if it exists, for every anchor. Chain score for every pair of anchors are derived from gap between anchors on the reference, gap between anchors on the query, overlap between anchors and average length of anchors as shown in Fig. 2b (adopted from Kalikar et al.^22^ and shown here for clarity). Optimal chain score generation is the most time consuming sub-task in chaining and is sequential within a read. For every anchor in a read, `mm2` proceeds sequentially through all the predecessors to generate chain scores and to find the optimal chain score as shown in Fig. 2a. However, `mm2` has a speed heuristic based on MAX_SKIP parameter which breaks out of the sequential predecessor check if a better scoring predecessor is not found beyond a certain number of total attempts (MAX_SKIP number of attempts) for any anchor. Prior works^22, 23^ have shown that removing this speed heuristic (by setting MAX_SKIP to infinity or INF) enables intra-read or more specifically intra-range parallelism (parallelizing the chain score generation with respect to all predecessors for any given current anchor) in chaining and also improves the mapping accuracy.

**Figure 2.**
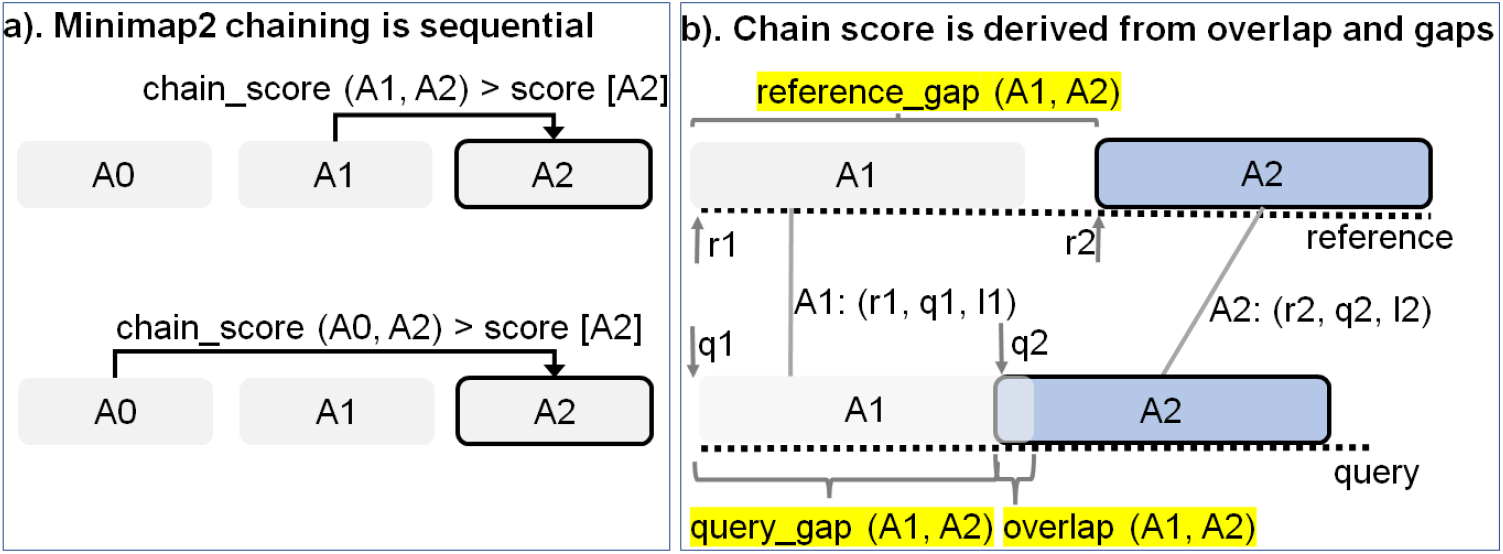
Chaining explained. (a) In Minimap2, every current anchor (A2 (r2, q2, l2) in this case) attempts to sequentially chain its predecessors within a pre-calculated predecessor range. If the chaining score with a predecessor is greater than the score value stored at current anchor A2, the new chain score and index of the predecessor is updated at A2 (in the direction of the arrow). (b) The chain score with a predecessor is computed from anchor gap cost (evaluated as a function of reference_gap, query_gap and average length of all anchors) and overlap cost.

The third sub-task identifies the maximum of scores at start and end of every chain per anchor and is sequential for every read. Predecessor range selection, chain score generation and finding maximum of scores at start and end of chain takes most of the time (97.42%) in chaining.

Backtracking and primary chain identification together takes only 2.58% of chaining time. Backtracking extends every anchor repeatedly to its best predecessor and ensures no anchor is used in more than one chain. Primary chain identification finds primary and secondary chains based on overlaps and estimates a mapping quality for each primary chain based on an empirical formula.

### Minimap2 profile

We profiled a single threaded CPU execution of `mm2` on randomly sub-sampled 100K reads of ONT and PacBio HiFi on an Intel Cascade Lake core and observed different profiles as previously noted^22^. Fig. 3 shows that for ONT, chaining is the bottleneck while alignment is the bottleneck for PacBio. Further, the percentage of time spent in chaining for ONT reads longer than 100Kb is as high as ∼68%. When the workload is normalized for the number of bases aligned, we also see that the long noisy ONT reads takes longer than PacBio HiFi on an average to align a base.

**Figure 3.**
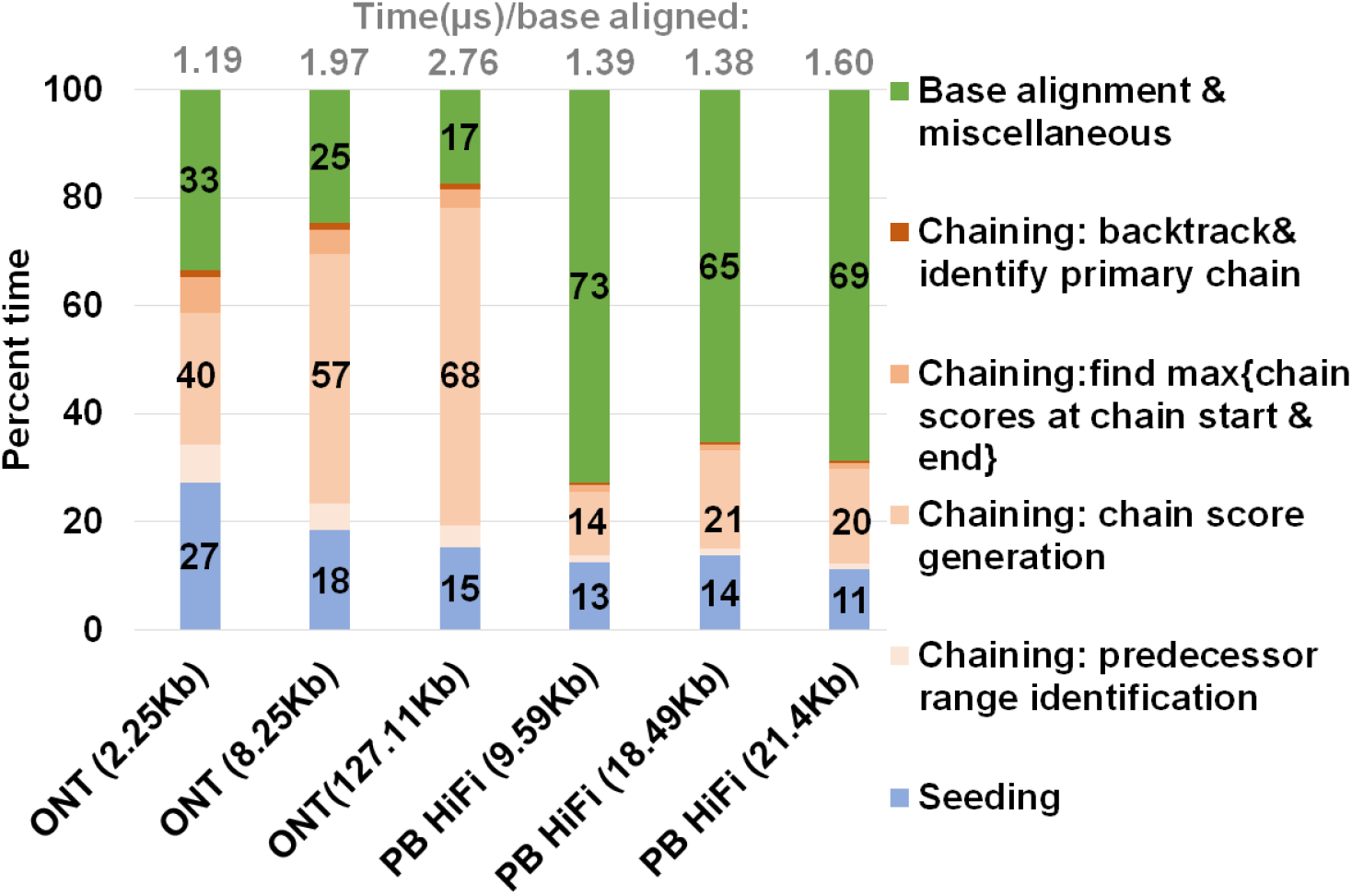
Summary of approximate time spent in seed-chain-align. `mm2` takes longer to map long noisy ONT reads and spends a greater percent of total mapping time in chaining. X-axis shows the sequencing technology with mean read length of each sets of 100K randomly sub-sampled reads.

Let us consider the randomly sub-sampled ONT dataset with 100K reads of mean read length 8.25Kb (second bar from the left in Fig. 3). This sub-set dataset is representative of the 60X HG002 dataset with N50 as 44Kb^24^ (N50 is an average read length metric used in genome assembly). Optimal chain score generation and finding the maximum of scores at start and end of a chain contribute to a significant part of the time spent in ONT chaining (90.9%). The other contributors to chaining are relatively smaller: predecessor range identification (6.6% of chaining), and backtracking and primary chain identification (2.5% of chaining).

Irregularity of workload (ONT reads vary in read lengths — a few hundred to a million bases), memory accesses, computation, and control flow associated with `mm2` makes accelerating it a difficult task. Further, `mm2` does not have any intra-read parallelism in chaining. Optimal chain score generation and finding the maximum of scores at start and end of a chain (which contribute to a total of 90.9% of the time in chaining) are implemented sequentially in `mm2`.

### Prior Work

There have only been a few prior works^22, 23, 25, 26^ which try to improve the performance of (accelerate) `mm2`. Zeni et al.^25^ and Feng et al.^26^ accelerate the base-level alignment step which is no longer the dominant bottleneck as reads have grown longer in length. Guo et al.^23^ and Kalikar et al.^22^ remove the MAX_SKIP heuristic for speed in `mm2` in order to extract intra-range parallelism and parallelizes chain score generation for each anchor (MAX_SKIP is set to INF). While Guo et al.^23^ correctly identifies chaining as the bottleneck for longer reads, introduces the concept of forward transforming the chaining algorithm and accelerates it on GPU and Field Programmable Gate Array (FPGA), this work fails to guarantee output equivalency to `mm2` with MAX_SKIP set to INF. We find that it misaligns (produces mismatched primary alignments) ∼7% of the reads with lengths above N50 while also failing to align ∼2% of those reads from our ONT 60X HG002 dataset. This decrease in mapping accuracy is mainly because Guo et al. follows a static predecessor range selection unlike the dynamic selection in `mm2` and also because the chaining score update rules are not modified accordingly with the transform.

`mm2-fast`^22^ is the most recent prior work in accelerating `mm2` and accelerates all three steps in `mm2` utilizing Single Instruction Multiple Data (SIMD processes multiple data with a single instruction) CPUs. While `mm2-fast` parallelizes chain score generation, we identify certain sections of chaining which are not parallelized. We profiled `mm2`-fast on the 100K sub-sampled reads from ONT and find that 34.08% of the total time spent in doing chain score generation and finding the maximum of scores at start and end of chains is in sequential code and not parallelized. `mm2-fast` does not use SIMD lanes when predecessor range is less than or equal to 5, for finding maximum predecessor score index and finding maximum of scores at start and end of chains for every anchor. This motivates the need for a better parallelization scheme.

### Our Contributions

In this work, we optimized the dominant bottleneck of `mm2` in processing long noisy reads, chaining, on the GPU without compromising accuracy. We show `mm2-ax` has better speedup and speedup : costup compared to `mm2-fast`, a SIMD-vectorized version of Minimap2 on 30 Intel Cascade Lake cores. As discussed, `mm2` presents a difficult task to parallelize with sequential chaining step and irregular workloads, memory accesses, computation, and control flow. Prior efforts at accelerating chaining either produces alignments significantly deviant from `mm2`^23^ or still does some amount of sequential execution within chaining and can benefit from a better parallelization scheme^22^. Hence, we attempted to better utilize the inherent parallelism in chaining without compromising accuracy on GPUs which are becoming increasingly popular for genomics workflows. To this end, we forward transform the predecessor range calculation to successor range calculation so as not to lose mapping accuracy and also forward transform the optimal chain score generation to introduce intra-range parallelism. Forward transformed chaining eliminates the need to sequentially find the maximum of all chain scores from all the SIMD lanes and instead enables better utilization of Single Instruction Multi-Threaded (SIMT is similar to SIMD but on a GPU) parallelization scheme on a GPU. Additionally, we also benefit from inter-read parallelism by concurrently processing multiple reads on the large number of Streaming Multiprocessors (SMs) on the GPU.

We designed a heterogeneous system where the bottleneck step, chaining, is sped up on the GPU while seeding and base-level alignment happens on the CPU. We exploit the low memory footprint of `mm2` and trade-off memory for performance via better occupancy of the GPU resources by the highly irregular workload in `mm2` chaining. Minimal branch divergence, coalesced global memory accesses and better spatial data locality are some of the optimizations.

We compare our accelerated minimap2 (`mm2-ax`) on GPU to SIMD-vectorized `mm2-fast` on CPU. Our evaluation metrics include accuracy, speedup, and speedup : costup. We show that `mm2-ax` produces 100% identical alignments to `mm2-fast` (same accuracy as `mm2` with MAX_SKIP set to INF) and delivers 5.41 - 2.57X speedup and 4.07 - 1.93X speedup : costup with respect to `mm2-fast` on ONT 60X HG002 dataset.

## Methods

### Parallelizing chaining: forward loop transformation

Chaining in `mm2` identifies optimal collinear ordered subsets of anchors from the input sorted list of anchors. `mm2` does a sequential pass over all the predecessors and does sequential score comparisons to identify the best scoring predecessor for every anchor. The exact chaining algorithm used in `mm2` is not parallelizable and hence, `mm2` is only able to utilize inter-read parallelism. Prior works^22, 23^ have shown that removing the speed heuristic in chaining by setting MAX_SKIP to INF enables intra-range parallelism (parallel chain score generation for all predecessors for any given anchor, i.e, parallelizing the inner for loop in Algorithm 1) and improves mapping accuracy. However, the total amount of work to be performed per anchor increases. We apply the same configuration in `mm2-ax`.

We find that ∼34% of the run time in `mm2-fast`’s optimal chain score generation and finding maximum of scores at start and end of all chains is spent sequentially. Chain score generation when the predecessor range is lesser than or equal to 5, finding maximum chaining score from among the 16 vector lanes and finding maximum of scores at start and end of all chains are all performed sequentially. In order to make better use of intra-range parallelism in chaining, we forward transform predecessor range selection (Fig. 4) and optimal chain score generation (Fig. 5 and Algorithm 1). This saves us the sequential passes which `mm2-fast` does to find the maximum chaining score.

**Figure 4.**
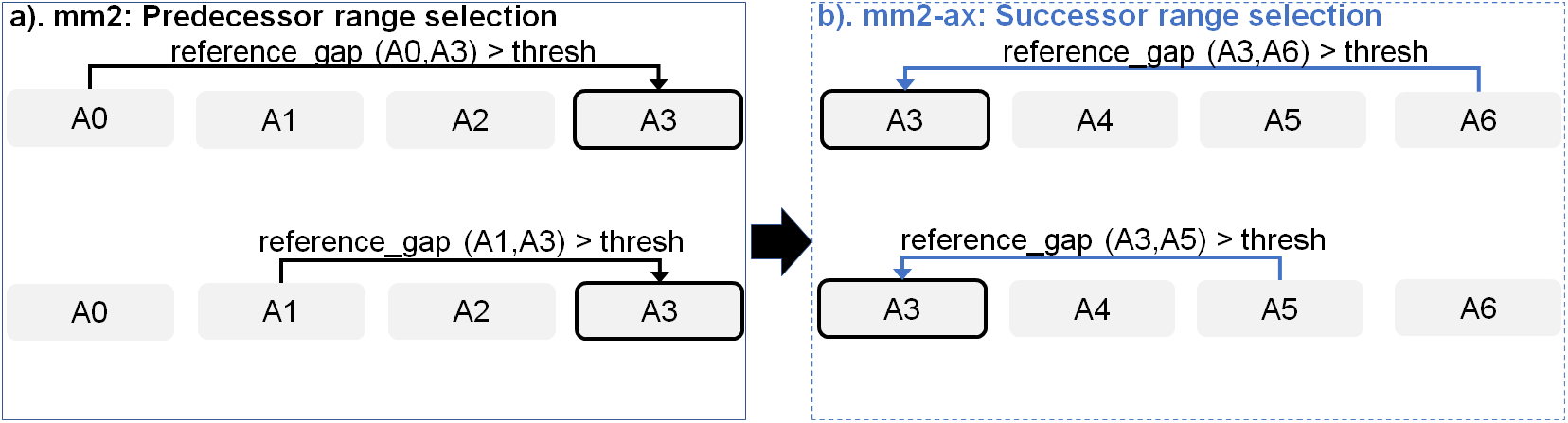
Forward transforming predecessor range selection to successor range selection: The cell with solid black outline represents the current anchor for which predecessor/successor range calculation is performed. The arrow starts from the predecessor/successor and points to the current anchor A3 whose range is updated sequentially.

**Figure 5.**
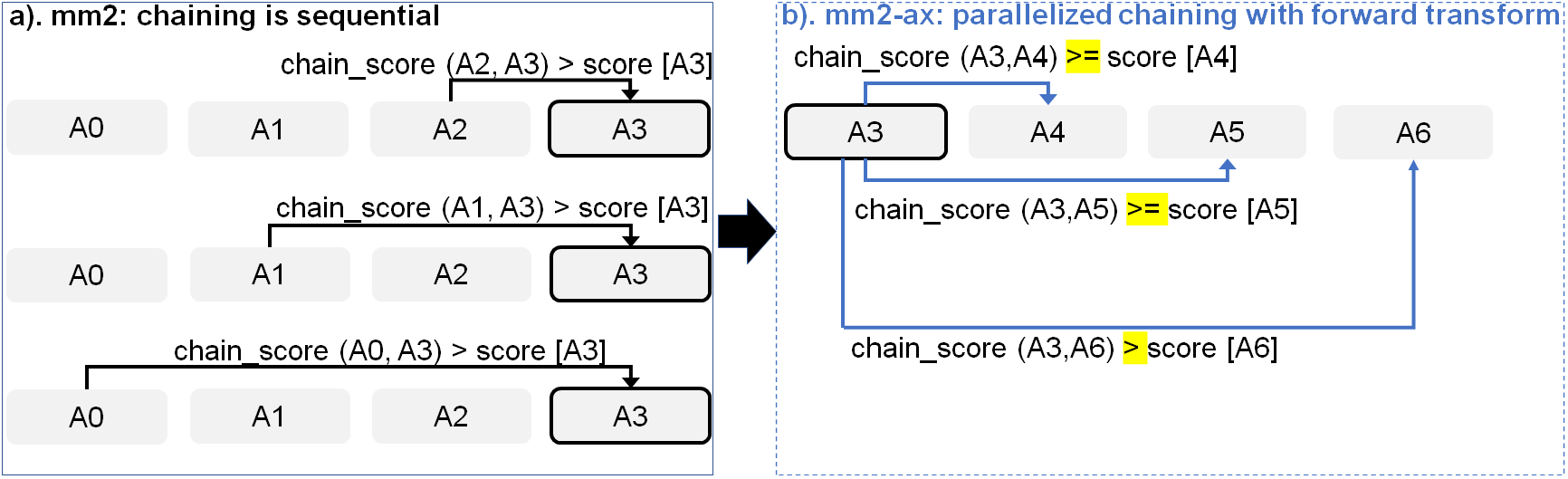
Parallelizing Minimap2’s chain score generation (shown in a) by forward transformation (shown in b). Additionally, we retain mapping accuracy by modifying the score comparison check (*>* to *>*=) with all anchors except the immediate neighbor to enable farther anchors to take precedence over neighboring anchors to be forward chained.

In this context, forward transformation refers to changing the order of computation to parallely evaluate successor anchors instead of iterating through predecessor anchors. This enables us to perform chain score generation and update in parallel as shown in Fig. 5. Although the forward transformation of optimal chain score generation is first introduced by Guo et al.^23^, in order to retain mapping accuracy, we implement two novel modifications. First, we calculate dynamic successor range instead of a static range of 64 for every anchor prior to chaining. We efficiently implement the successor range calculation with few iterations based on insights from cumulative distribution function of predecessor ranges for all anchors (discussed later in Fig. 6b). Secondly, the chain score update policy is modified from *>* to ≥ (except for the immediately neighboring anchor) for the forward traversal as shown in Fig. 5b. This ensures that farther anchors get precedence over nearer ones for forward chaining.

**Figure 6.**
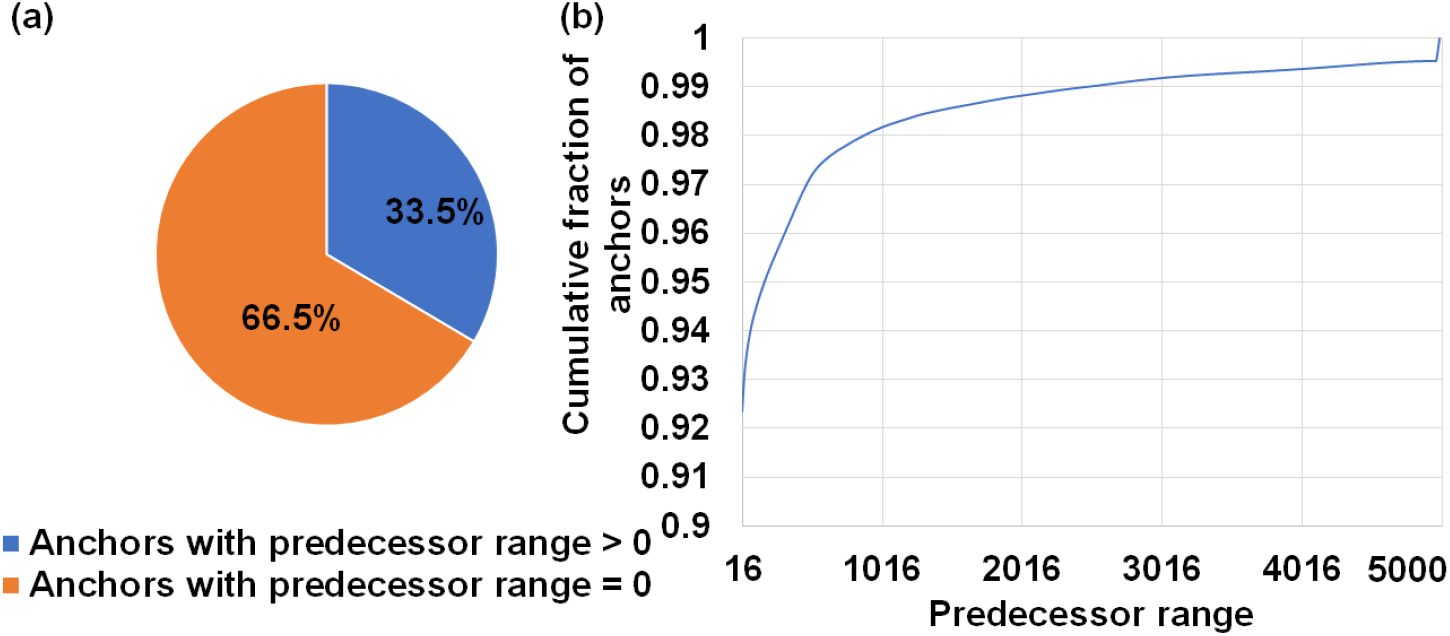
Workload is sparse and irregular. (*a*) ∼67% of anchors fed to the chaining step do not start a chain. (b) Predecessor range is less than or equal to 16 for ∼92% of all anchors and goes as high as 5000 only for a small fraction of total anchors.

#### Algorithm 1 Forward transformed chaining in *mm2-ax*

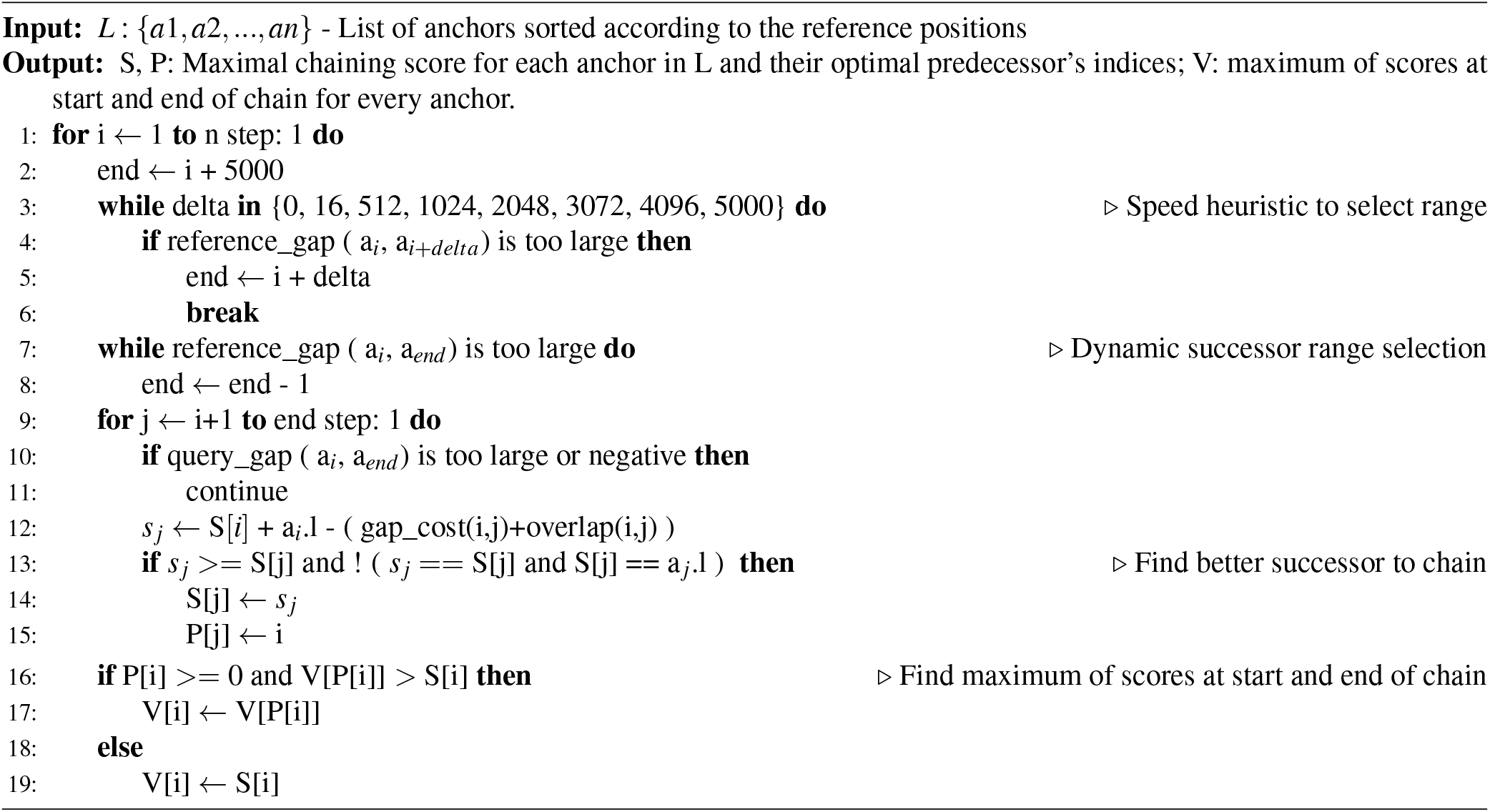

### Heterogeneous system design

`mm2-ax` is a heterogeneous design (uses specialized compute cores, GPUs in this case) which performs seeding and successor range identification on the CPU and efficiently implements optimal chain score generation and finding maximum of scores at start and end of chain on the GPU. The output scores and optimal successor index arrays from chaining are returned to the host CPU for backtracking. From `mm2`’s profile in Fig. 3, optimal chain score generation and finding maximum of scores at start and end of chain contribute 90.6% of chaining time and is accelerated on the GPU. Seeding, successor range identification or forward transformed predecessor range identification (6.6% of chaining), backtracking and primary chain identification (2.5% of chaining) and base-level alignment are performed on the CPU.

Further, the heterogeneous design also helps us better balance the workload and reduce resource idling on the GPU, as discussed in below sections.

### GPU occupancy: Condensed workload vector and workload balancing

We find that ∼67% of the input anchors do not start a chain and this contributes to the sparsity of the successor range vector which is to be input to optimal chain score generation. In order to better occupy the GPU resources with the irregular workload, we perform successor range identification (steps 3-8 in Algorithm 1) on the CPU to convert this sparse input vector of successor ranges which defines the workload in chain core generation, into a condensed one with non-zero successor ranges. This incurs a GPU and host memory trade-off for better performance by ensuring GPU threads do not idle on anchors with a successor range of zero. Further, the compute overhead on the host CPU from successor range identification is minimized by implementing a speed heuristic (steps 3-6 in Algorithm 1) to reduce the number of iterations in identifying the successor range for every anchor. This is based on the observation that ∼67% of the anchors on an average have a predecessor/successor range of zero and ∼93% have a range lesser than or equal to 16.

Further, we also implement a series of additional measures to ensure better GPU occupancy, as we realize that this is one of the most important problems^27^ while dealing with ONT reads of variable lengths and predecessor ranges. To ensure GPU occupancy from workload balancing, we bin and batch reads of similar lengths together onto the GPU. For example, reads of length 2Kb-3Kb, 3Kb-4Kb and 4Kb-5Kb are binned together. For smaller read lengths (*<*= 10Kbp), we define each concurrently launched workload at a coarser grain, i.e, as many reads as it takes to concurrently occupy all the SMs on the

GPU. For longer reads, we observe that this does not yield the optimal performance because long reads present a case of highly imbalanced workloads as reads are more variable in length and any SM which may finish early remains unused. For example, in 50-150Kb range, reads are highly varying in read lengths, and it is difficult to find multiple reads within 1Kb variance in lengths. Hence, for longer reads we keep bin ranges wider: 45-50Kb, 50-100Kb and 100-150Kb. For longer reads, we follow a two-fold strategy for higher GPU occupancy. First, we define fine-grained workloads, i.e, with only as many reads as it takes to occupy an entire SM. Second, we always follow up very-long read bin workloads with fine-grained workloads of shorter read lengths (2Kb). This twofold strategy helps better balance highly imbalanced workloads of very long reads.

For better GPU occupancy, we also launch multiple concurrent GPU kernels (functions) using CUDA streams (GPU work queues). As soon as a hardware resource gets free on the GPU, the scheduler executes the next kernel. Additionally, each Streaming Multiprocessor (SM) concurrently processes multiple reads.

Data transfer between the CPU and GPU are overlapped with compute on the GPU by issuing asynchronous memory copies on CUDA streams. We also benefit from the higher bandwidth of HBM2 and the eight copy engines on A100.

### Inter-read and intra-range parallelism

A server-class GPU like NVIDIA A100 has 108 SMs. The key to high performance on the GPU is to ensure that all the SMs always have useful work to do and there are sufficient Single Instruction Multi-Threaded (SIMT) warps/sub-warps (groups of threads) concurrently on the GPU to hide the relatively higher global memory access latencies (i.e, ensure higher warp occupancy). While we utilize only inter-read parallelism (concurrently processing multiple reads per SM) for finding the maximum of scores at start and end of chain, we utilize both inter-read and intra-range parallelism (via forward transformation) for the optimal chain score generation. Intra-range parallelism comes from concurrent warps (sets of 32 parallel threads) performing chain score generation in parallel for all successors within the successor range of a given anchor.

The next anchor attempts to chain only after it’s previous anchor’s optimal chain score generation step is completed. To this end, we have a thread synchronization barrier (__syncthread()) waiting on all the threads to finish chain score generation and update for all the successors of a given anchor. Please note that Guo et al.^23^ uses more synchronization barriers (six of them) in the chain score generation kernel. However, we only need one as we reduce the number of points of branch divergence by combining multiple condition checks together.

### Data Locality

We observe optimal benefits from optimizing for better spatial data locality rather than temporal locality. Temporal cache locality refers to re-use of data in cache, while spatial cache locality refers to use of data from adjacent storage locations. For example, we pre-fetch data for a group of successors per current anchor in each concurrently processed read into L1 cache for better performance from improved spatial data locality. We use the PTX instruction prefetch_global_l1 to prefetch the successor anchor’s inputs (query and reference coordinates) and chaining output (score and parent values) from global memory to L1 cache in a coalesced fashion for every set of 32 successors per anchor.

While Guo et al.^23^ attempted to exploit temporal locality from using shared memory (memory shared between parallel threads of a read) with a static successor range, this approach does not prove beneficial with a dynamic successor range because of limited scope for any benefits from temporal data locality. Frequent cache misses due to different successor ranges lead to data transfer latency from shared memory to registers, adding up to outweighing any benefit from using shared memory at all. We therefore use more registers per GPU thread instead of utilizing shared memory.

We also coalesce global memory reads and writes for successor anchors to reduce the total number of transactions to high-latency global memory.

### Minimal branch divergence

Conditional branches are kept to a minimum in our implementation by combining conditions when successors are not updated after score generation. This helps reduce branch divergence, which affects performance on the GPU. There are only two conditional blocks for every read that is processed within the chain score generation kernel (one for score generation and the other for update, as seen in Algorithm 1). On the other hand, Guo et al.^23^ has nine conditional blocks evaluated per read.

Further, we utilize CUDA’s warp-synchronized integer intrinsics to efficiently perform operations like logarithm and absolute differences. __clz() lets us efficiently calculate logarithm during the chain score generation step from counting the leading zeros and subtracting this count from the number of bits in int32 datatype (32). __sad() enables us to efficiently compute the overlap cost from the absolute difference of query_gap and reference_gap (shown in Fig. 2b).

## Implementation

### Experimental Setup

Minimap2 (`mm2`) is a fast evolving software with 7 new releases on the master branch and 2 new branches incorporating `mm2-fast` in the year 2021 alone. We decided to accelerate Minimap2 v2.17 which is used in Oxford Nanopore’s variant calling pipeline with Medaka^9^. Kalikar et al.^22^ has accelerated Minimap2 versions v2.18 and v2.22. `mm2` v2.18 and v2.17 produce equivalent results in chaining and are, hence, comparable.

We demonstrate the benefits of our chaining optimizations on a server class NVIDIA A100 GPU. Our evaluations are performed on Google Cloud Platform (GCP). Speedup is normalized to a costup factor to evaluate a speedup : costup metric in order to take into account the usually higher GPU costs on the cloud. Our costup factor on GCP is 1.33X, but this would be lower if one were to use Amazon Web Services (AWS). `mm2-ax` is evaluated on a single GCP instance of a2-highgpu-1g with 85GB of host memory and one NVIDIA A100 GPU of 40GB memory. We compare `mm2-ax` to the SIMD accelerated `mm2-fast` (`fast-contrib` branch of `mm2`) on a single GCP c2-standard-60 instance (30 AVX-512 vectorized Intel Cascade Lake cores and 240GB memory).

We use NVIDIA Nsight Compute^28^ for profiling GPU events and Nsight Systems^29^ for visualization of concurrent GPU events. Seqtk^30^ is used for random sub-sampling of DNA sequences. We used Perf^31^ for profiling `mm2` on the CPU.

To ensure better GPU resource utilization with nanopore reads of varying length (few hundred to a million bases), we bin reads based on read lengths before batching their sorted anchors onto the GPU for chaining. For example, reads of length 1-2 Kilobases (Kb) go to the same bin, reads of length 2-3Kb go to the same bin etc. However, for longer read lengths, we bin 50K-100K, 100K-150K etc. because it is relatively harder to find reads closer in read lengths.

The reads within a bin could still present an unbalanced workload as the predecessor ranges of every anchor is different. This binning may be done very efficiently during basecalling as the basecaller has access to read lengths, and hence the overhead introduced is negligible. We also try to fit in as many reads as possible on to the GPU’s DRAM for every read bin. We measure the compute time for optimal chain score generation and sub-task to find maximum of scores at start and end of chain on the GPU and compare it to that of the SIMD baseline to evaluate SpeedUp metric (compute time taken by CPU baseline `mm2-fast` divided by time taken by `mm2-ax` on GPU). We then divide this with 1.33X to normalize for cost and calculate the speedup : costup metric. The overhead presented by successor range selection over predecessor range selection on the host CPU is very negligible (*<* 2.8% of total CPU time) and is, hence, not considered for our analysis. Further, it is worthwhile to note that successor range identification can outperform predecessor range identification using SIMD vectorization on the host CPU as our forward transform essentially makes successor range identification parallelizable.

Further, we also evaluate mapping accuracy of `mm2-ax` vs `mm2-fast` (or `mm2` with MAX_SKIP set to INF). Mapping accuracy is defined as the number of reads from `mm2-ax` producing bit-exact chains to `mm2-fast`. If any of the 12 fields in `mm2-ax`’s Pairwise Alignment Format (PAF) formatted output differs from that of `mm2`’s in the primary alignments, we treat the read as misaligned. The datasets we use are publicly available^32–35^ — HG002 genome sequenced by ONT PromethION with 60X coverage and 15Kb and 20Kb PacBio HiFi reads with 34X coverage.

### Optimal GPU conifgurations

Of the two chaining sub-tasks offloaded to the GPU, chain score generation takes approximately greater than 95% of the time on the GPU. Hence, we discuss how we performed design-space exploration to identify the optimal GPU kernel launch parameters for this sub-task. Kernel launch parameters refer to a predefined configuration with which a kernel or function may be executed on the GPU. In this context, we can define the chain score generation kernel launch parameters as a 3-tuple (thread blocks per SM, number of concurrent reads processed per block in an SM, number of parallel threads per read). In this context, thread blocks are groups of parallel threads within an SM which may or may not be processing the same read.

We find the register requirement per thread on an NVIDIA A100 GPU to figure out the achievable upper bound of GPU kernel launch parameters on the A100 GPU. Using NVIDIA’s Nsight Compute Profiler, we profiled `mm2-ax` and observed that we require 53 registers per thread for the optimal chain score generation kernel, and this observation helps provide an upper bound on the maximum number of parallel threads that can be launched on the SM in our case. From Fig. 6b, one may try to fit more concurrent reads with 16 or 32 threads per read, but it is observed that this configuration hurts spatial cache locality across reads and is hence, not beneficial. The optimal configuration is observed to be in the direction of higher concurrent reads per SM instead of per thread bock and towards more threads allocated per read for chaining. This is because having more threads per read enables better spatial data locality in L1 cache through larger coalesced global memory accesses. We find that (9 thread blocks per SM, 1 concurrent read per thread block, 128 parallel threads per read) is the best performing kernel configuration. This is followed by (3, 3, 128) and (1, 4, 256).

From Fig. 6a, we observe that ∼67% of anchors fed to the chaining step do not start a chain. This observation helps us to ensure better arithmetic intensity (more computations per byte of data fetched from high latency global memory). In this regard, we perform successor range identification on the host CPU and condense the sparse vector of successor ranges to a dense one with non-zero successor range before offloading the chain score generation sub-task to the GPU. Further, Fig. 6b informed us to efficiently implement successor range identification. 67% of the anchors have predecessor ranges equal to zero, and greater than 92% have predecessor ranges less than or equal to 16. We use this information to efficiently implement successor range selection by reducing the number of total iterations.

We did a design space exploration to identify the granularity of dispatching work to the GPU for optimal performance. While concurrently dispatching coarse grained workloads (each workload is defined with as many reads as it takes to fill all the 108 SMs on A100) yielded the best results for reads smaller than or equal in length to 10Kb, coarse grained workloads do not work well for longer reads as any SM that finishes early may idle. Concurrently dispatching fine-grained workloads (each workload is sized with as many as reads as it takes to fill only a single SM) yielded the best performance for reads longer than 10Kb. Concurrent fine-grained dispatches with each workload sized to fill 2 and 4 SMs closely competes but gives slightly lower benefits. Additionally, a fine-grained workload for longer reads is always followed by fine-grained workload of smaller reads (we choose 2Kb) to yield a better performance. This is because multiple fine-grained workloads of smaller reads can better load balance without adding a significant tail to the critical path.

## Results

`mm2-ax` demonstrates a 5.41 - 2.57X speedup and 4.07 - 1.93X speedup : costup over SIMD-vectorized `mm2-fast` baseline as shown in Fig. 7a. It is observed that coarse-grained load dispatching to the GPU is better for read lengths smaller than 10Kb while fine-grained load dispatching to each SM is better for longer reads. In Fig.7b we show the chaining performance gain factor without including the data transfer related costs (memory allocation, asynchronous memory copies on CUDA streams and data serialization).

**Figure 7.**
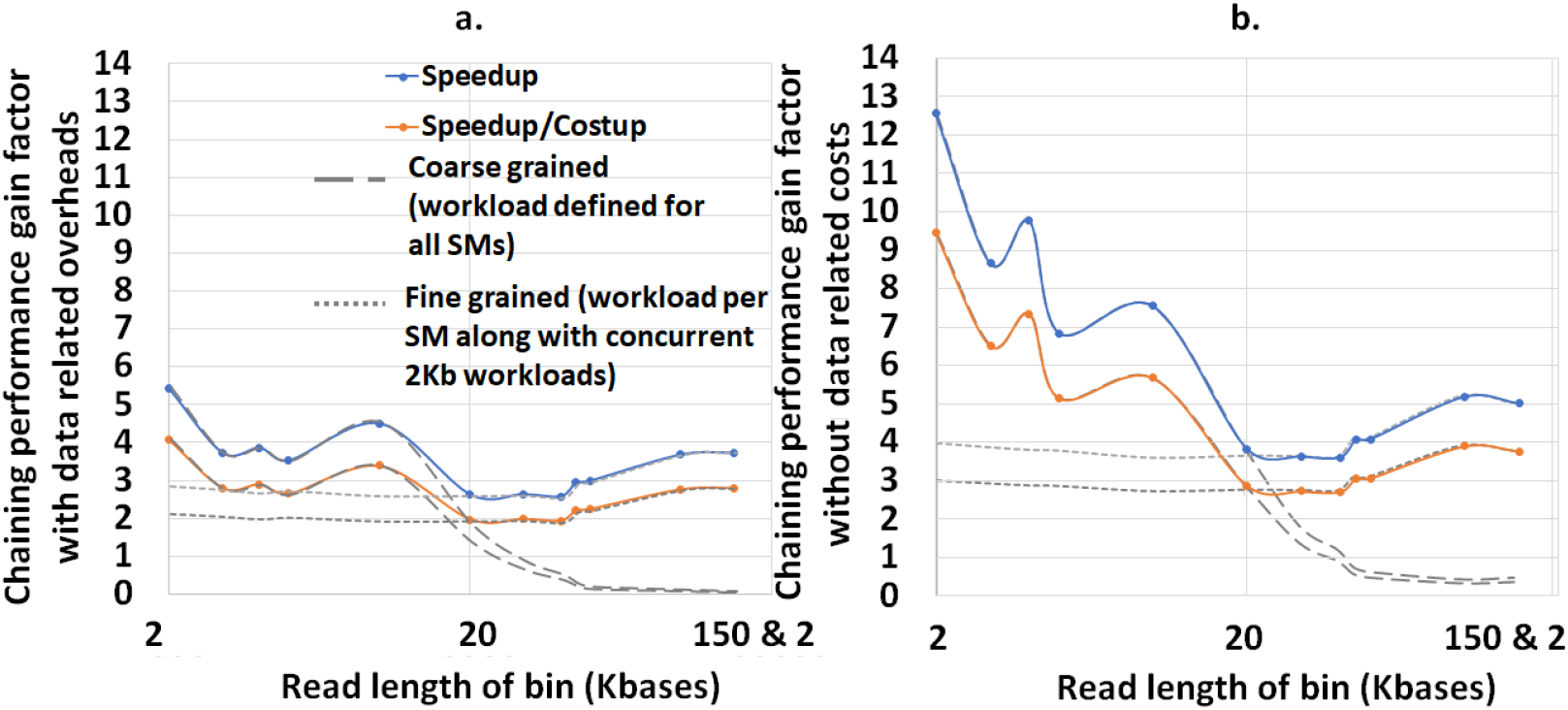
(a) mm2-ax yields 5.41 - 2.57X speedup and 4.07 - 1.93X speedup : costup over SIMD-vectorized *mm2-fast* baseline. (b) The chaining performance across various read lengths may be further improved by ∼1.3-2.3X if we can engineer to hide the data transfer related costs.

Additionally, we also evaluate the mapping accuracy of `mm2-ax` with respect to `mm2-fast` on the complete 60X HG002 ONT dataset and we observe that all the output chains match in all the first 12 fields of the output Paiwise Mapping Format (PAF) file. As `mm2-ax` is limited by the DRAM capacity of the GPU, we use minimap2 modified with our forward transformed chaining logic on the CPU to evaluate accuracy on the very large 60X HG002 dataset. We then compare and validate the output PAF file to the outputs of both `mm2-fast` and `mm2` modified with MAX_SKIP set to INFINITY.

## Discussion

We only discuss the profile of the most time-consuming kernel of `mm2-ax` on the GPU, optimal chain score generation. Here we present the profile of optimal chain score generation kernel on the 2Kb bin of reads. From Fig. 8a, we see that the chain score generation kernel on the GPU is memory bound. Unless we increase the arithmetic intensity of the kernel, we cannot transform it to a compute bound kernel. From Fig. 8b, we observe that high register usage per thread limits the theoretical number of active warps per SM on the GPU. The higher the warp occupancy, the better the kernel is in hiding the relatively longer global memory access latency on the GPU. The achieved number of active warps per SM is 12 (33% of the theoretical maximum). One way to ensure there are enough warps on the SM is by integrating asynchronous FIFOs at the input and output of chaining to better manage input and output out of the GPU. One may also try to better hide the data transfer related costs to gain further improve in performance.

**Figure 8.**
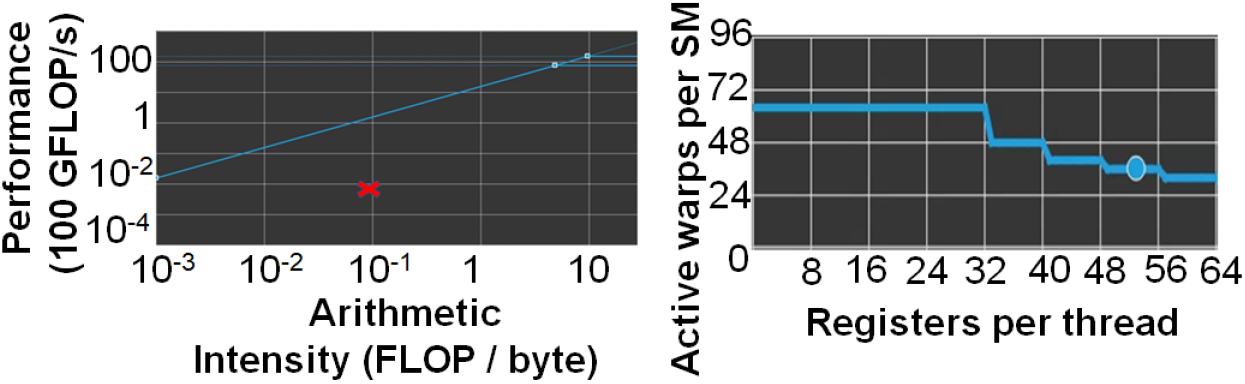
mm2-ax is memory bound. (a)Roofline Plot: Chain score generation is memory bound. Operating point is shown in a red cross mark. (b) Theoretical warp occupancy on the GPU is bounded by the number of registers used by each thread.

Further, it may be noted that one may further improve performance by making the range of read lengths that go into a bin even smaller. In our case, the HG002 dataset presented reads in 50-150Kb range to be highly varying in read lengths and hence, we defined long read bins with higher variance in read lengths. It is also worthwhile to consider porting the entire `mm2` software to the GPU as most long read sequencing workflows are now shifting to GPUs.

## Conclusions

Long read sequencing workflows on GPUs are becoming increasingly popular for healthcare and genomics applications like Precision Medicine and microbiome abundance estimation. We identify chaining as the bottleneck in the state-of-the-art aligner used for long read mapping and alignment. Chaining constitutes as high as ∼67% of the total alignment time in long noisy ONT reads of lengths greater than 100Kb. We address this problem with `mm2-ax` (minimap2-accelerated), a heterogeneous system for accelerating the chaining step of minimap2. We implement various optimizations to ensure better occupancy and workload balancing on the GPU. Some key optimizations include forward transformed chaining for better intra-read parallelism, trading-off host and GPU memory for better performance on the GPU, better spatial data locality and minimal branch divergence. We show `mm2-ax` on an NVIDIA A100 GPU improves the chaining step with 5.41 - 2.57X speedup and 4.07 – 1.93X speedup : costup over the fastest version of minimap2, `mm2-fast`, benchmarked on a single Google Cloud Platform instance of 30 AVX-512 vectorized cores (Intel Cascade Lake).

## Acknowledgements

Development of `mm2-ax` was supported by NVIDIA Corporation and University of Michigan Ann Arbor (via D. Dan and Betty Kahn foundation grants). Additionally, we would like to thank Oded Green, Lotfi Slim, Harry Clifford, Mehrzad Samadi and Ajay Tirumala from NVIDIA Corporation for their helpful suggestions and advice on efficiently using NVIDIA GPUs.

## Author contributions statement

H.S. performed the analysis, design, and implementation of `mm2-ax`. M.M. recommended various performance optimizations and CUDA best practices for implementing `mm2-ax`. E.D. and V.I. helped better understand the `mm2` chaining algorithm and prior work. J.I. and S.N. led the collaborative effort and helped design the optimization strategy for `mm2-ax`. All authors reviewed the manuscript.

## Data availability statement

Datasets are publicly available with CC0 license from Human Pangenome Reference Consortium: https://github.com/human-pangenomics/HG002_Data_Freeze_v1.0^32–36^.

## Additional information

### Software Availability

`mm2-ax` is currently closed source. However, a docker image for `mm2-ax` and instructions on using the image with singularity is made available on GitHub: https://github.com/hsadasivan/mm2-ax. Zenodo DOI: https://doi.org/10.5281/zenodo.6374533.

### Competing interests

The authors declare that they have no competing interests.

## Notes

https://s3-us-west-2.amazonaws.com/human-pangenomics/index.html?prefix=NHGRI_UCSC_panel/HG002/hpp_HG002_NA24385_son_v1/nanopore/downsampled/standard_unsheared

https://github.com/human-pangenomics/HG002_Data_Freeze_v1.0

### Competing Interest Statement

The authors have declared no competing interest.

### Summary of Updates

Revised with updated evaluations with data transfer overheads

